## Supplemental Information for "Integrating When and What Information in the Left Parietal Lobe Allows Language Rule Generalization"

**Supporting Information**

Orpella J^1, 2, 3, 4^*, Ripollés P^4^*, Ruzzoli M^5^, Amengual JL^6^, Callejas A^1,7^, Martinez A^1, 2, 3^, Soto-Faraco S^5, 8^, de Diego-Balaguer R^1, 2, 3, 8^.

^1^Cognition and Brain Plasticity Unit, IDIBELL, L’Hospitalet de Llobregat, Spain

^2^Dept of Cognition Development and Educational Psychology, University of Barcelona, Barcelona, Spain

^3^Institute of Neuroscience, University of Barcelona, Barcelona, Spain

^4^Dept of Psychology, New York University, New York, USA

^5^Center for Brain and Cognition, Departament de Tecnologies de la Informació i les Comunicacions, Universitat Pompeu Fabra, Barcelona, Spain

^6^Centre de Neuroscience Cognitive Marc Jeannerod, CNRS UMR 5229, Université Claude Bernard Lyon I, Bron, France

^7^Departamento de Psicología Experimental, Facultad de Psicología y Centro de Investigación Mente, Cerebro y Comportamiento, Universidad de Granada, Spain.

^8^ICREA, Barcelona, Spain

* Equal contribution

**Corresponding author**

**Ruth de Diego-Balaguer**

**Faculty of Psychology**

**Pg. Vall d’Hebron 171**

**08035 Barcelona**

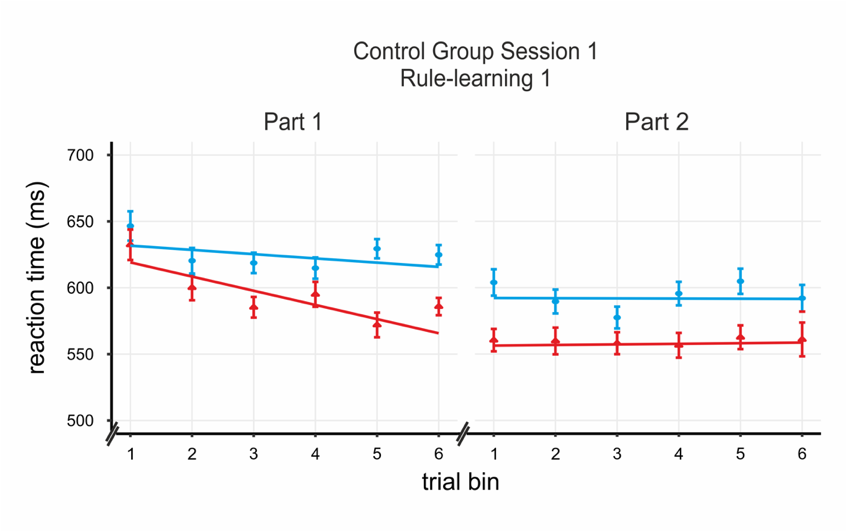

**Figure S1. Control group incidental rule-learning task results for Session 1.** Slopes for rule and no-rule blocks over task repetitions derived from the mixed model analysis. The Control Group showed the expected transition from a significant learning slope in Part 1 (ßdiff = -0.8, *t* = -3.1, *p* < 0.002) to a non-significant learning slope (ßdiff = 0.06, *t* = 0.252, *p* > 0.8) with a significant rule effect in Part 2 (*t*(30) = 4.49, *p* < 0.001) and Part 1 (*t*(29) = 3.6, *p* < 0.002). Actual data shown averaged into 6 trial bins per part with the SEM for display purposes.

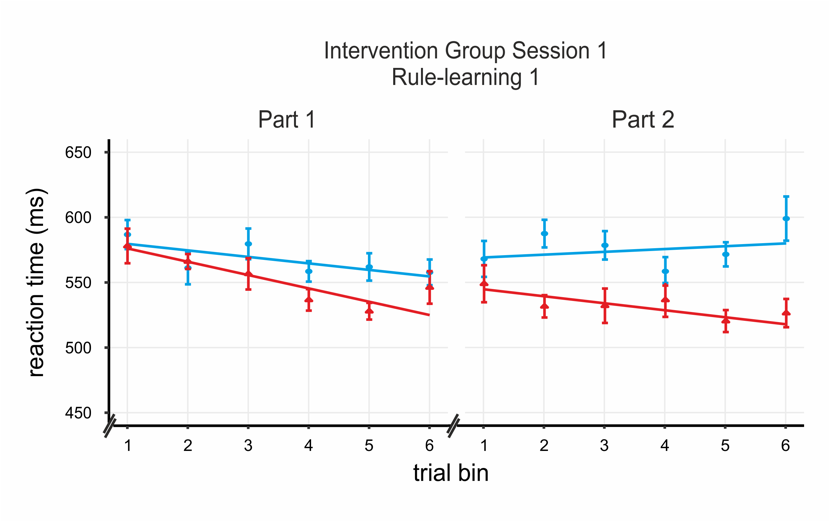

**Figure S2. Intervention group incidental rule-learning task results for Session 1.** Slopes for rule and no-rule blocks over task repetitions derived from the mixed model analysis. the Intervention Group –perhaps owing to the smaller sample and/or scanner effects– appeared to comprise slower learners who still showed a significant learning slope during fMRI phase (Part 2: ßdiff = -0.8, *t* = -2.63, *p* < 0.01; Part 1: ßdiff = -0.56, *t* = -2.13, *p* < 0.034), as well as the expected rule effects (Part 1: *t*(16) = 2.19, *p* < 0.044; Part 2: *t*(19) = 3.08, *p* < 0.007). Actual data shown averaged into 6 trial bins per part with the SEM for display purposes.

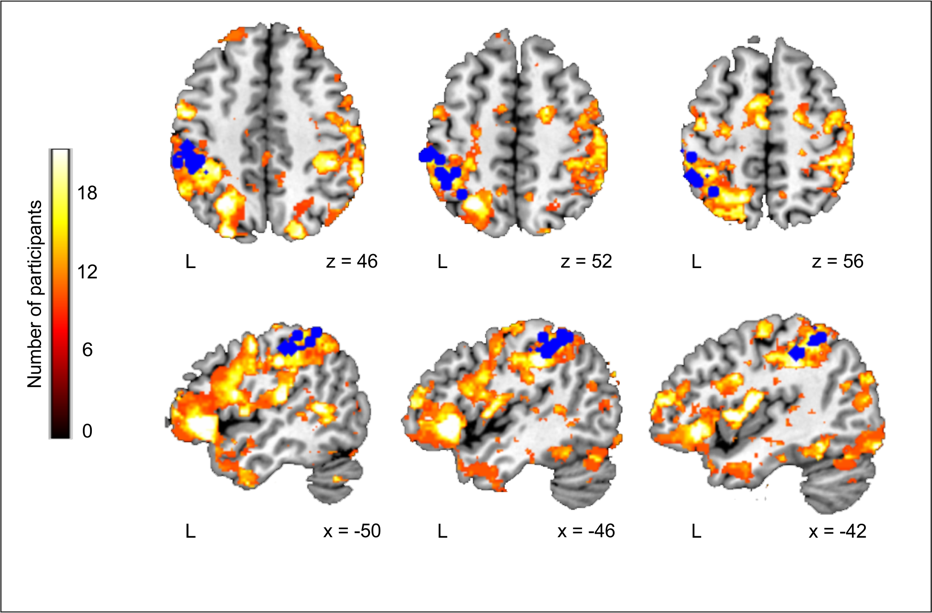

**Figure S3. Individual fMRI enhanced activity during later stages of rule learning and rTMS stimulation sites.** In red-yellow, overlap of individual masks for each participant's activation pattern. Only voxels in which at least 10 participants showed individual fMRI enhanced activity during Rule learning are shown. In blue, the sites for rTMS stimulation for all participants is shown (for clarity purposes, 4 mm spheres were created around the stimulation centers). Neurological convention is used with MNI coordinates shown at the bottom right of each slice.

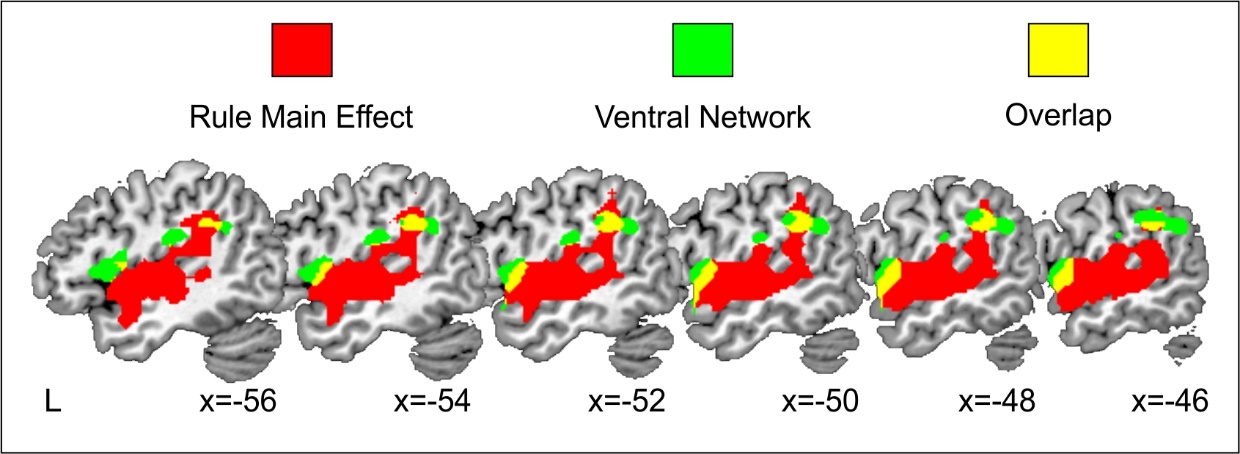

**Fig. S4. Overlap (yellow) between the ventral network correlating with Statistical Learning (green) and the main rule effect (Rule vs. Baseline; in green).** In left ventral frontal and parietal regions there is an overlap between the contrast showing regions in which the BOLD signal significantly co-varies with the measure of statistical learning (Learning Slope) and the contrast showing the brain regions in which activity increases during Rule blocks: the more activity in these regions during rule blocks, the greater (i.e. more negative) the slope. Only significant results (*p* < 0.05 FWE-corrected at the cluster level, with an additional *p* < 0.001 at the voxel level and 50 voxels of cluster extent) are shown over a canonical template with MNI coordinates on the bottom right of each slice. L, Left Hemisphere.

**Table S1. Whole brain fMRI activity related to individual differences in rule slopes.** Group-level fMRI local maxima for areas correlating with negative (i.e. RTs are still decreasing) rule slopes during the rule blocks in the fMRI phase (see also red-yellow regions in **Fig. 4A**, main text). Results are reported at a *p* < 0.05 corrected threshold at the cluster level with 50 voxels of cluster extent, with an additional uncorrected *p* < 0.005 threshold at the voxel level. MNI coordinates were used. BA, Brodmann Area.

| Anatomical area | Coordinates | Cluster Size | t-value |
| --- | --- | --- | --- |
| Left Inferior Parietal Lobe (BA 40)  Left Postcentral gyrus (BA 2) | -54 -40 32 | 924 | 6.89 |
| Left Inferior Frontal Gyrus (BA 44,45)  Left Insula (BA 13) | -54 12 2 | 936 | 6.77 |
| Right Inferior Frontal Gyrus (BA 44,45)  Right Insula (BA 13) | 44 16 6 | 545 | 5.17 |

**Table S2. Whole brain fMRI activity related to individual differences in the rule effect increment for Part 2.** Group-level fMRI local maxima for areas correlating with rule effect increments in Part 2 for the rule vs. no-rule contrast during the fMRI phase (see also red-yellow regions in **Fig. 4B**). Results are reported at a *p* < 0.05 corrected threshold at the cluster level with 50 voxels of cluster extent, with an additional uncorrected *p* < 0.005 threshold at the voxel level. MNI coordinates were used. BA, Brodmann Area.

| Anatomical area | Coordinates | Cluster Size | t-value |
| --- | --- | --- | --- |
| Left Mid/Sup. Frontal Gyrus (BA 8,9)  Left Precentral Gyrus (BA 6) | -40 26 36 | 1650 | 7.12 |
| Right Pallidum  Bilateral Caudate | 16 0 -2 | 655 | 6.09 |
| Bilateral Precuneus (BA 7, 5)  Bilateral Post/Mid. Cingulate Gyrus (BA 24, 31)  Bilateral Inf./Sup. Parietal Gyrus (BA 40, 7)  Bilateral Postcentral Gyrus (BA 3) | -14 -62 32 | 4277 | 5.90 |
| Left Postcentral Gyrus (BA 2,3)  Left Inf. Parietal Gyrus (BA 40) | -34 -22 48 | 894 | 5.85 |
| Right Mid./Sup. Frontal Gyrus (BA 9,10) | 32 34 28 | 721 | 5.48 |

**Table S3.** Coordinates for parietal stimulation sites.

| Subject | Coordinates | | |
| --- | --- | --- | --- |
| 1 | -58 | -32 | 50 |
| 2 | -44 | -46 | 52 |
| 3 | -56 | -30 | 50 |
| 4 | -50 | -26 | 46 |
| 5 | -48 | -42 | 50 |
| 6 | -54 | -30 | 52 |
| 7 | -44 | -36 | 44 |
| 8 | -55 | -54 | 38 |
| 9 | -38 | -46 | 60 |
| 10 | -48 | -46 | 56 |
| 11 | -42 | -34 | 44 |
| 12 | -56 | -36 | 44 |
| 13 | -46 | -38 | 46 |
| 14 | -44 | -50 | 54 |
| 15 | -48 | -34 | 54 |
| 16 | -50 | -30 | 46 |
| 17 | -34 | -56 | 54 |
| 18 | -58 | -48 | 40 |
| 19 | -38 | -42 | 50 |
| 20 | -46 | -36 | 46 |

**Offline recognition test**

*Methods*

Following each online learning phase, participants’ knowledge of the rules was additionally assessed via a recognition test. Participants were presented with phrases that conformed to the rules in half of the trials and phrases that violated them in the other half. Incorrect sentences were, in half of the cases, violations of the dependency using A and C elements from different rule structures in their correct position but violating the specific A_C *dependency* (i.e. A1xC2, A2xC1), and, in the other half, they were *order* violations, whereby A and C elements swapped positions within the phrase albeit maintaining their specific dependency (i.e. C1xA1 and C2xA2). Correct sentences were in half of the trials the same sentences previously presented and in the other half they were new sentences that contained an x from the pool of x exposed in the language but not previously combined with the specific A1_C1 or A2_C2 dependency. The complete offline test thus comprised a total of 48 test phrases (24 per rule dependency). Participants were instructed to discriminate between phrases that could and could not belong to the previously heard language by pressing the corresponding button. Button-response relation was counterbalanced across participants. No limits on response time were set in the baseline and rTMS tests, though participants were instructed to respond quickly after the whole phrase was heard. A new phrase was delivered immediately after a participant’s response. In the fMRI scanner, a response time maximum threshold of 1500 ms was set, along with a jittered interval between 1000 and 3000 ms before the start of the next trial.

Participants’ ability to discriminate rule items from violations was assessed by transforming to *d* prime scores (*d′*) accuracy responses. For each participant, the proportion of hits (i.e. *yes* responses to rule phrases) and false alarms (i.e. *yes* responses to violations) were used to calculate the *d′* score after taking care of hit and false alarm rates of zero or one (49). We computed three different *d’* scores by calculating the false alarms using: i) both order and dependency trials (*d’_All_)*, ii) order violations (*d’_Ord_*), and iii) dependency violations (*d’_Dep_*). These scores were then submitted to one-sample and paired *t*-tests to test for statistical significance (*d′* = 0 corresponding to no discrimination). In order to assess the effects of the rTMS lPL intervention, *d’* in this condition was compared to *d’* scores in the rTMS POz intervention in the same subjects. In addition, *d’* under the rTMS lPL effects was also compared to *d’* in the Control group as a between subjects comparison.

*Results*

Participants from both groups were able to significantly discriminate sentences that followed the learned dependencies from all violations in all phases (Session 1 Part 1: Intervention group, *d’All* = 0.34 ± 0.44, *t*(16) = 3.185, *p* < 0.01, d_Cohen_ = 0.773, Control group, *d’All* = 0.42 ± 0.46, *t*(31) = 5.13, *p* < 0.001, d_Cohen_ = 0.907; Session 1 Part 2: Intervention group, *d’All* = 0.55 ± 0.80, *t*(18) = 3.03, *p* < 0.01, d_Cohen_ = 0.696, Control group, *d’All* = 0.69 ± 0.92, *t*(31) = 4.24, *p* < 0.001, d_Cohen_ = 0.749; rTMS lPL: Intervention group, *d’All* = 0.54 ± 0.74, *t*(19) = 3.25, *p* < 0.004, d_Cohen_ = 0.729; rTMS POz: *d’All* = 0.55 ± 0.71, *t*(19) = 3.45, *p* < 0.003, d_Cohen_ = 0.773; Control group Session 2: *d’All* = 0.85 ± 0.80, *t*(31) = 5.96, *p* < 0.001, d_Cohen_ = 1.045). When specifically looking at the effects of the rTMS intervention on Session 2 we observed it had no effect on the *d’* discrimination scores. Performance was comparable in the rTMS lPL and the rTMS POz interventions in the within subjects comparison (*d’All: t*(19) = 0.11, *p* = 0.912, d_Cohen_ = 0.025) and there were no significant differences for the comparison of the rTMS lPL and the rTMS POz conditions to the performance of the Control group in the between subjects comparisons (respectively, *d’All: t*(50) = 1.39, *p* =0.170, d_Cohen_ = 0.397 and *t*(50) = 1.33, *p* = 0.189, d_Cohen_ = 0.379).

**Table S4.** Results for the d’ calculated with the order or the dependency as false alarms

| **Violation** |  | **Session** |  |  |
| --- | --- | --- | --- | --- |
|  | **Session 1** | | **Session 2** | |
|  | **Part 1** | **Part 2 (fMRI)** | **rTMS POz** | **rTMS IPL** |
| **d’_Ord_ CxA**  **Intervention Group** | d’_Ord_ = 0.75 ± 0.78  *t*(16) = 3.94  *p* < 0.001  d_Cohen_ = 0.95 | d’_Ord_ = 1.15 ± 1.01  *t*(18) = 5.01  *p* < 0.001  d_Cohen_ = 1.14 | d’_Ord_ = 1.10 ± 0.88  *t*(19) = 5.55  *p* < 0.001  d_Cohen_ = 1.24 | d’_Ord_ = 0.98 ± 1.15  *t*(19) = 3.83  *p* < 0.001  d_Cohen_ = 0.85 |
| **d’_Dep_ A1xC2**  **Intervention Group** | d’_Dep_ =-0.04 ± 0.51  *t*(16) = -0.30  *p* = 0.76  d_Cohen_ = -0.07 | d’_Dep_ = 0.06 ± 0.89 *t*(18) =0.32  *p* = 0.74  d_Cohen_ = 0.07 | d’_Dep_ = 0.09 ± 0.86 *t*(19) = 0.50  *p* = 0.62  d_Cohen_ = 0.11 | d’_Dep_ = 0.21 ± 0.79 *t*(19) = 1.18  *p* = 0.25  d_Cohen_ = 0.26 |
|  | **Part 1** | **Part 2** |  | |
| **d’_Ord_ CxA**  **Control Group** | d’_Ord_ = 0.75 ± 0.78  *t*(31) = 5.01  *p* < 0.001  d_Cohen_ = 0.88 | d’_Ord_= 1.05 ± 1.19  *t*(31) = 4.98  *p* < 0.001  d_Cohen_ = 0.88 | d’_Ord_= 1.25 ± 1.01  *t*(31) = 6.98  *p* < 0.001  d_Cohen_ = 1.236 | |
| **d’_Dep_ A1xC2**  **Control Group** | d’_Dep_ = 0.08 ± 0.47  *t*(31) = 1.02  *p* = 0.315  d_Cohen_ = 0.180 | d’_Dep_ = 0.37 ± 0.97 *t*(31) = 2.19  *p* < 0.04  d_Cohen_ = 0.387 | d’_Dep_ = 0.50 ± 1.08  *t*(31) = 2.62  *p* < 0.02  d_Cohen_ = 0.463 | |

When the *d’* was calculated for each type of violation separately, we observed that in all sessions and groups, participants were sensitive to order (CxA) indicating they learned the position of the dependencies (see Table S4). Indeed, learning of position was maintained despite the rTMS intervention on Session 2 and was observed in all conditions with no differences between them (rTMS POz to rTMS lPL within subjects comparison: *d’*_Ord:_*: t*(19) = 0.57, *p* = 0.57, d_Cohen_ = 0.129; rTMS POz to Control between subjects comparison: *d’*_Ord_*:*t(50) = 0.56, p = 0.57, d_Cohen_ = 0.161; rTMS lPL to Control between subjects comparison: *d’*_Ord_*: t*(50) = 0.95, *p* =0.34, d_Cohen_ = 0.271). However, only the Control group was sensitive to the dependency violations (A1xC2; see Table S4) pointing that only this group, which was tested three times with no interference in any of the three languages learned, was able to benefit from the repetitive testing to extract this more detailed knowledge of the dependencies in Session 2 (i.e. generalization session). Nevertheless, no strong conclusions can be drawn from this result since there were no differences in Session 2 between rTMS POz and rTMS lPL in the within subjects comparisons (d’_Dep_*: t*(19) = 0.59, *p* = 0.56, d_Cohen_ = 0.133) but also no significant differences in the between subjects comparisons (rTMS POz to Control group: *d’*_Dep_: t(50) = 0.98, p = 0.330, d_Cohen_ = 0.280; rTMS lPL to Control group: *d’*_Dep_*: t*(50) = 1.41, *p* =0.160, d_Cohen_ = 0.403.

**Table S5.** Stimuli set for the rule-learning task

| X stimuli L3 | A_C structures: | | X stimuli L2: | A_C structures: | | X stimuli L1: | A_C structures: | |
| --- | --- | --- | --- | --- | --- | --- | --- | --- |
| baki | A1 | lexa | bade | A1 | kote | bidu | A1 | tagi |
| bosi | C1 | kudo | bilu | C1 | naxu | defa | C1 | sipa |
| dela | A2 | bedu | dimu | A2 | dufa | fako | A2 | pine |
| farre | C2 | moga | duga | C2 | tulo | foli | C2 | ladu |
| gadu |  | | fiko |  | | gosa |  | |
| gumi |  |  | fusi |  |  | gupe |  |  |
| kinu |  |  | gabe |  |  | katu |  |  |
| lemo |  |  | gexo |  |  | lasu |  |  |
| line |  |  | kire |  |  | lofa |  |  |
| mexa |  |  | ledo |  |  | male |  |  |
| mogi |  |  | lifa |  |  | medi |  |  |
| nifa |  |  | loke |  |  | mego |  |  |
| nosa |  |  | mefi |  |  | nigo |  |  |
| nuxo |  |  | nipa |  |  | nuso |  |  |
| pate |  |  | nuta |  |  | pebo |  |  |
| pula |  |  | pafi |  |  | pote |  |  |
| rrugo |  |  | pugo |  |  | pume |  |  |
| sato |  |  | roku |  |  | rrosu |  |  |
| sofu |  |  | sipu |  |  | sapu |  |  |
| teku |  |  | some |  |  | supa |  |  |
| tipe |  |  | teTo |  |  | tadi |  |  |
| todi |  |  | Tune |  |  | Tilu |  |  |
| xapu |  |  | xabu |  |  | Toba |  |  |
| xebo |  |  | xagi |  |  | turre |  |  |
